# Delineation of post-phloem assimilate transport pathway into developing caryopsis of *Brachypodium distachyon*

**DOI:** 10.1101/718569

**Authors:** Charles Ugochukwu Solomon, Sinead Drea

## Abstract

Assimilates stored in mature cereal grains are mobilized from source tissues and transported towards developing grains through the vascular bundle. Due to the lack of direct vascular connection between maternal grain vascular bundle and filial tissues, post-phloem transportation of assimilates into grain endosperm relies on transfer cells that lie between the grain vascular bundle and the endosperm. Here, we propose Caryopsis Endosperm Assimilate Acquisition Route (CEAAR) models that describes the exact path of assimilate import into caryopsis endosperms. Using fluorescent tracer dyes we also delineated the route of assimilate delivery into Brachypodium distachyon endosperm and classified it as ventral circuitous (vc-CEAAR), an assimilate import model also found in rice. Furthermore, we report a detailed anatomical study of post-phloem assimilate transport pathway in developing grains of *Brachypodium distachyon*. Our results highlight major anatomical similarities and differences between the grain post-phloem transfer cells of Brachypodium and those of crop species such as rice, wheat, and barley relevant to post-phloem assimilate transport.

**Highlights:**

- Based on existing work, we propose Caryopsis Endosperm Assimilate Acquisition Route (CEAAR) models, that describes the exact path of assimilate import into caryopsis endosperms.
- The structure of the post-phloem transfer cells of *Brachypodium distachyon* mirrors temperate and tropical cereals.
- Assimilate delivery into *Brachypodium distachyon* endosperm is identical to assimilate import into rice endosperm.

## Introduction

Cereals are major source of food and feed for humans and their livestock. The value of cereals lies in the dry storage capacity of their endosperm. An endosperm is a sink tissue, therefore assimilates stored in the endosperm originate from source tissues in other parts of the plant. During grain filling, assimilates are translocated from source tissues towards the endosperm through the vascular bundle. Because of the absence of vascular continuity between maternal source tissues and filial sink tissues, assimilates are unloaded from the phloem and traverse post-phloem transfer cells on their journey to the endosperm. Thus, the post-phloem transport pathway serves as a bridge connecting maternal supply to filial sink of cereal caryopsis. Based on studies of assimilate movement into developing filial tissues of cereals like rice, maize, sorghum, barley and wheat, three models for post-phloem transport pathway into caryopses endosperms can be deduced. These models, which we collectively termed Caryopsis Endosperm Assimilate Acquisition Route (CEAAR) models are; basal direct (bd), ventral circuitous (vc), ventral direct (vd) and (Fig. 1). Barley and wheat represent the first model; vd-CEAAR. In these species, assimilates are also delivered by a ventrally located grain vascular tissue that run the length of the grain. On exit from the phloem, assimilates symplastically cross chalazal cells, nucellar projection cell sand accumulate temporally in an apoplastic space called the endosperm cavity. Specialized endosperm transfer cells (also called modified aleurone cells) handles the importation of assimilates from the endosperm cavity into the endosperm (Fig. 1a) (Cochrane, 1985;Cochrane and Duffus, 1980; Donovan et al., 1983; Frazier and Appalanaidu, 1965; H. L.Wang, C. E. Offler, et al., 1995; H. L. Wang, J. W. Patrick, et al., 1995; Zheng and Wang, 2011).

**Fig. 1:**
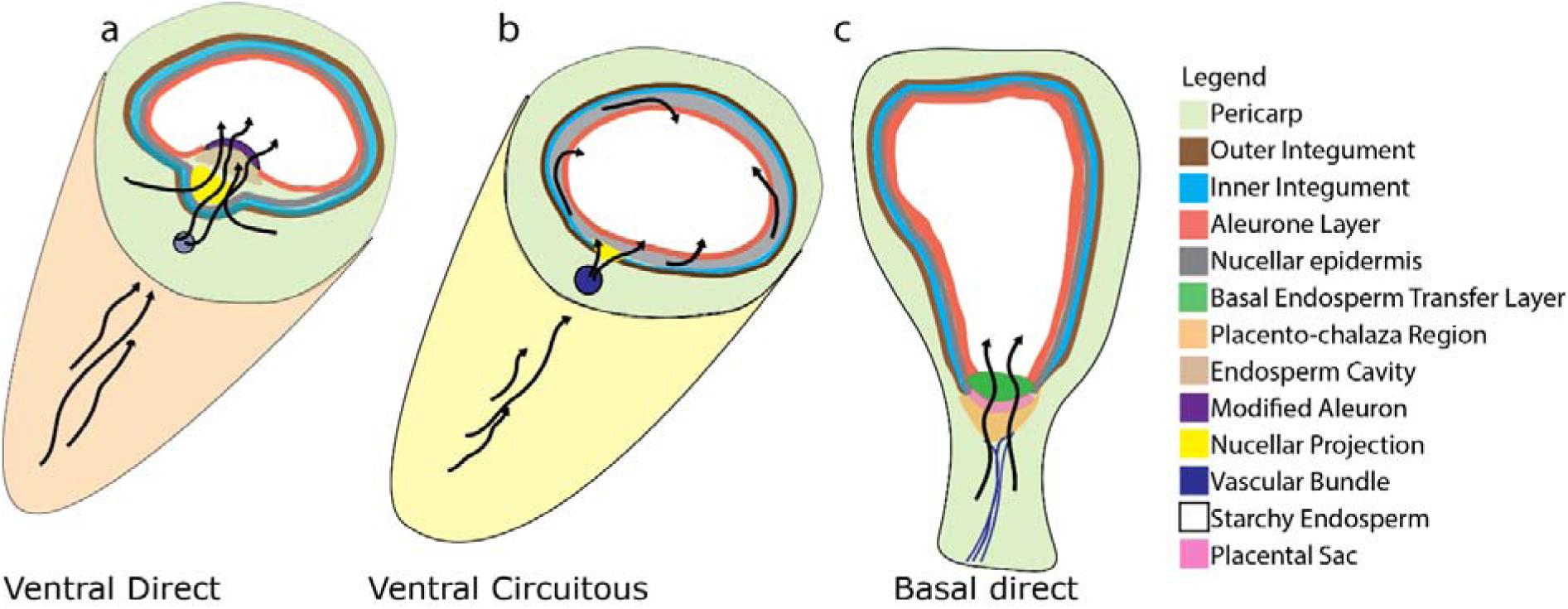
Caryopsis Endosperm Assimilate Acquisition Route (CEAAR) models. The models illustrate how resources are mobilized into the endosperm of grass caryopsis.

An example of vc-CEAAR is found in rice, where assimilates are delivered by the ovular vascular tissue that runs the entire length of the ventral surface of developing rice grains. Assimilates symplastically traverse the chalazal zone, a remnant of nucellar and move into nucellar epidermis cells. Within the nucellar epidermis, they move circumferentially around the developing endosperm and eventually cross the plasma membrane of the nucellar epidermis cell into an apoplastic space adjoining the aleurone cells. From this space they are imported into aleurone cells and further into starchy endosperm cells (Fig. 1b) (Ellis and Chaffey, 1987; Krishnan and Dayanandan, 2003; Oparka and Gates, 1981, 1982; Wu et al., 2016). The third model, bd-CEAAR, is seen in maize and sorghum. Here the vascular trace ends at the base of the grain. Assimilates headed for storage in the endosperm are released from the phloem sieve tubes, from there they cross the chalazal strip and remnant nucellar cells. In sorghum and some varieties of maize, assimilates accumulate in the placental sac which serves as a transient sink. Basal Endosperm transfer layer (BETL) cells are responsible for transport of assimilates from the placental sac into the developing endosperm (Fig. 1c) (Felker and Shannon, 1980; Maness and McBee, 1986; Thorne, 1985; Zheng et al., 2015).

The importance of post-phloem assimilate transfer cells for proper caryopses development is easily manifest when there is perturbation of normal gene regulation and expression in the transfer cells. For example, maize *MINIATURE1* (*MN1*) and rice *GRAIN INCOMPLETE FILLING* (*GIF1*) loci encodes cell wall invertases that are expressed in the transfer cells of maize and rice grains respectively. Null mutation of MN1 or mis-regulation of GIF1 results in smaller grains compared to wild type (Cheng et al., 1996; Miller and Chourey, 1992; Wang et al., 2008). Similarly in barley, mature grains of shrunken endosperm genetic (seg); seg1, seg3, seg6 and seg7 mutants are shrunken because post-phloem transfer cells collapse early compared to wild type, leading to premature cessation of grain filling (Bosnes et al., 1992; Felker et al., 1985, 1984, 1983). Furthermore, maize *ZmSWEET4c* and its rice ortholog *OsSWEET4* are hexose transporters expressed in post-phloem transfer cells. Their loss of function mutation results in incomplete grain filling (Sosso et al., 2015). In view of the importance of the expression of *GIF1, OsSWEET4* and *ZmSWEET4c* in the post-phloem transfer cells of rice and maize respectively, for proper grain filling, they no doubt contribute to large grain trait of these cereals and are therefore postulated to have been recruited during the domestication of these cereals (Doebley et al., 2006; Sosso et al., 2015; Wang et al., 2008).

*Brachypodium distachyon* (subsequently, Brachypodium) on the other hand is a wild grass that recently became a model system for temperate cereals. Brachypodium grains are comparable to barley and wheat in length, but are narrower in width and shallower in depth (Hands and Drea, 2012). Although, median transverse section profiles of Brachypodium grains are generally identical to barley, wheat and oat sections, Brachypodium grains have reduced nucellar projection and persistent nucellar epidermis (Kosina and Tomaszewska, 2016; Opanowicz et al., 2011). Notably, modified aleurone, critical for assimilate import into the endosperm of barley and wheat is absent in Brachypodium grains. This attribute suggests that the final step of assimilate import into Brachypodium endosperm employs a different route compared to wheat and barley. Hence, we speculated that assimilate movement into Brachypodium endosperm follows the vc-CEAAR model as found in rice (Hands et al., 2012). In this study, we used fluorescent dyes; 5-(6)-6 carboxyfluorescein diacetate (CFDA) and Lucifer Yellow (LW) to trace the path of assimilate movement into Brachypodium endosperm.

We confirm that the route of assimilate import into Brachypodium endosperm is similar to rice and can be classified as vc-CEAAR. In addition, we report detailed ultrastructural study of caryopsis post-phloem transfer cells of Brachypodium. Our results highlight the anatomical basis for the route of assimilate transport into Brachypodium endosperm. Structural similarities and differences between the caryopses post-phloem transfer cells of Brachypodium and major cereals relevant to assimilate import into the endosperm are discussed.

## Materials and Methods

### Growth of Brachypodium plants

Brachypodium (Bd 21) Grains, were imbibed on moist filter paper in a Petri dish and left at 5°C for two days to vernalize. They were transferred to room temperature (about 25-27°C) and left to germinate. After 7 days, the most virile seedlings were transferred to 9:1 Levington M2 Pot and Bedding Compost: Levington Fine Vermiculite mix (dejex.co.uk), in Vacapot™ 15 on plastic seed trays (www.plantcell.co.uk). They were grown in the greenhouse at 16hr daylight and 25°C temperature. The plants were regularly watered manually. Flowering spikes were tagged at anthesis and sampled at 5 days interval after anthesis until 30 days post anthesis (DPA).

### Fluorescence tracer experiments

Fluorescence dyes used are symplastic tracer; 0.01% 5-(6)-6-carboxyfluorescein diacetate (CFDA; Sigma) and apoplastic tracer; 1% Lucifer Yellow (LY; Sigma). Sterile distilled water served as control. Brachypodium grains were carefully detached from their rachis at 20 DPA and their lemma removed under a dissecting microscope. About 20 grains were incubated in 1 ml solution of either fluorescent dyes or water in 1.5 ml Eppendorf tubes at room temperature. Five grains were sampled from each treatment at 15, 30, 45 and 60min. This was followed by a vigorous rinse in 1 ml distilled water three times before the grains were transferred to 0.5 ml 50% glycerol. Free hand 80 – 150 μm transverse sections spanning the length of the grains were made with clean razor blades. The sections were mounted on 50% glycerol covered with cover slips and the edges sealed with nail varnish. Imaging was done with Nikon ECLIPSE 80i fluorescence microscope (Nikon, Japan), having an LED-based excitation source (CoolLED, presicExcite), using Nikon Plan Fluor 10x /0.30 DIC L/N1 objective lens. Fluorescence images were captured with a DS-QiMc cooled CCD camera (Nikon, Japan). Images were previewed, captured and saved using NIS-Elements Basic Research v3.0 software (Nikon, Japan) in JPEG2000 format.

### Brachypodium grain processing for light and transmission electron microscopy

Individual Brachypodium florets were tagged at anthesis and developing grains were sampled at five days interval until 30 days after anthesis (DPA), six developmental stages in total. At each interval, five grains were collected from different florets. 1 mm section was cut from the mid grain region of each grain, using a dissecting microscope. Sections were fixed in 2.5% glutaraldehyde in 0.1M sodium Cacodylate buffer pH 7.4, for 3 days at 4°C with constant gentle agitation and then washed in 0.1M Sodium Cacodylate buffer. After further fixation in 1% aqueous osmium tetroxide and dehydration in series of increasing ethanol concentrations followed by propylene oxide, the seed sections were embedded in Spurr’s hard resin and polymerised for 16 hours at 60°C.

### Light microscopy

Grain sections for light microscopy were cut 400nm thick, stained with 0.01% toluidine blue, mounted in resin, then viewed and photographed with GX L3200B compound microscope (gtvision.co.uk) equipped with a CMEX-5000 USB2 Microscope camera and ImageFocus 4 software (euromex.com). Images were stored as TIFF files and annotated with Adobe Illustrator.

### Electron microscopy

70nm thick sections for transmission electron microscopy were cut using a Reichert Ultracut E ultra microtome. The sections were collected onto copper mesh grids, and stained with 2% aqueous uranyl acetate for 30 minutes followed by 5 minutes in lead citrate. Sections were viewed on a JEOL JEM-1400 TEM (www.jeolusa.com) with an accelerating voltage of 100kV. Digital images were collected with a Megaview III digital camera with iTEM software (emsis.eu) as TIFF files.

## Results

### Cellular structure of assimilate transfer route in Brachypodium grains Vascular bundle

Groups of sieve element – companion cells (se – cc) complex was observed on the ventral (adaxial) vascular bundle on all the grains sampled at all stages. The number of se – cc complex varied between grains at the same developmental stage but generally increased as the grain develops. Characteristic of monocots, the se – cc complexes were irregularly scattered between vascular parenchyma cells (Fig. 2b, d-g), except at 10 DPA when they seemingly formed a semi-circle facing towards the pigment strand (Fig. 2a, c). Mixtures of mature (empty lumen) and immature (lumen with remnants of cytoplasm) sieve elements cells were observed in all sample stages (Fig. 3a). The accompanying companion cells were not always easily identified. They were smaller and usually had denser cytoplasm compared to vascular parenchyma cells (Fig. 3a). Plasmodesmata connections were observed between sieve elements and their companion cells, as well as between sieve elements and vascular parenchyma cells. Similarly, there were plasmodesmata connections between companion cells and vascular parenchyma cells. Remarkably, we could not identify any xylem elements in sections from all the stages.

**Fig. 2:**
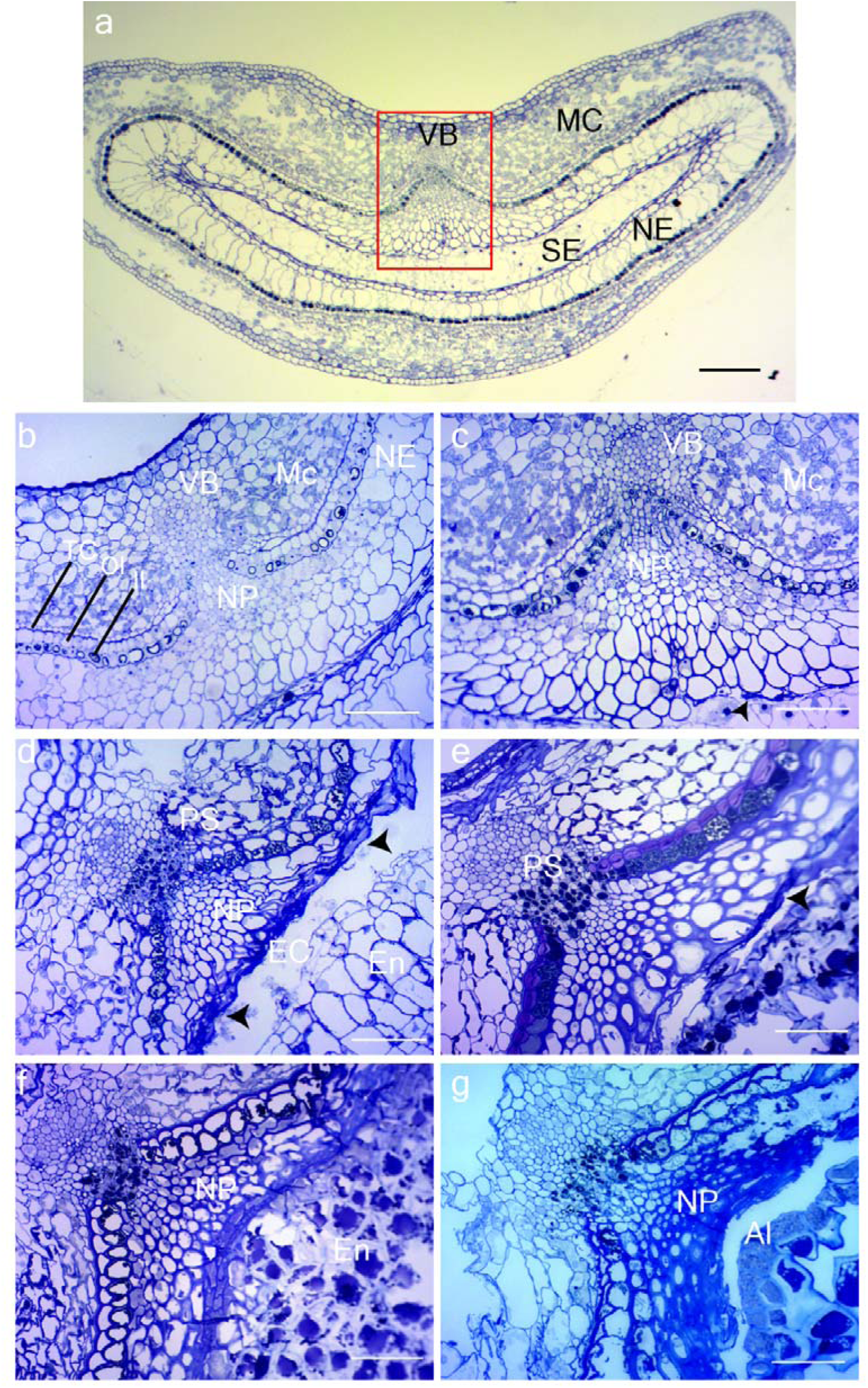
Development of Brachypodium distachyon grain post-phloem assimilate transport pathway. (a)Transverse section of a Brachypodium grain at 10 days post anthesis (DPA). Red square indicates area magnified in subsequent images. (b), (c), (d), (e), (f), and (g) are sections Brachypodium grain post-phloem assimilate transport route at 5, 10, 15, 20, 25, and 30 DPA respectively. Sieve element-companion cell complex were present in the vascular bundle all sections. There was progressive accumulation of dark deposits in the pigment strand as the grain develops. Cell death and disintegration of nucellar projection cells closest to the endosperm was observed from 10 DPA and is indicated with arrow heads. This created a transient endosperm cavity present at 15 and 20 DPA. The thickness of nucellar epidermis cell wall adjoining the endosperm increased as the grain developed. Bar = 0.1 mm. Al, aleurone cells; TC, tube cell; EC, endosperm cavity; II, inner integument; NE, Mc, mesocarp; nucellar epidermis; NP, nucellar projection; OI, outer integument; PS, pigment strand; En, endosperm; VB, vascular bundle.

**Fig. 3:**
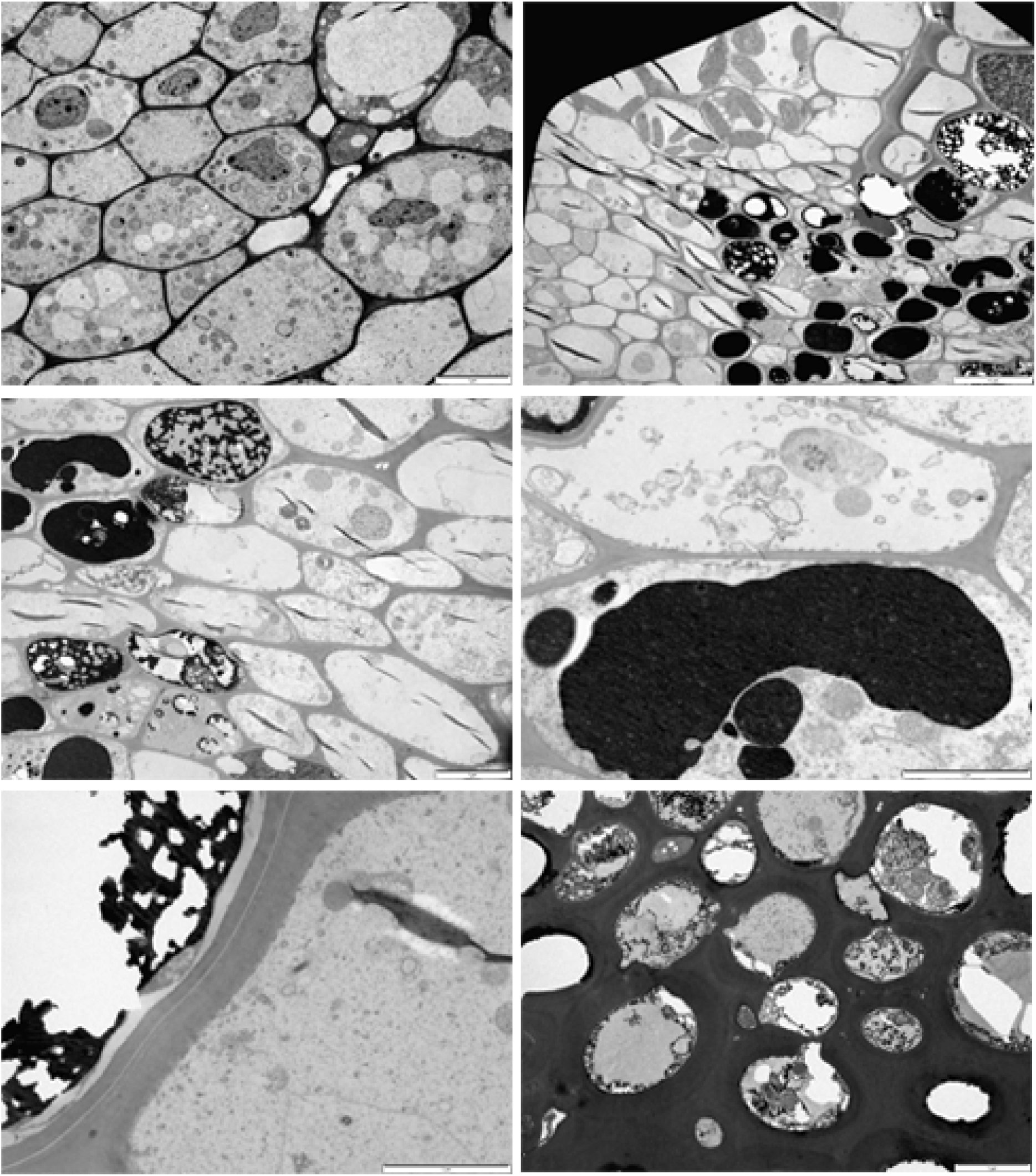
Transmission electron micrographs of transverse sections of developing Brachypodium grains. (a) Vasuclar bundle at 20 DPA. Note the prominent nucleus and presence of numerous mitochondria (M) in the dense cytoplasm of the vascular paranchyma (VP) cells. Enucleate seive elements (SE) are also present. Bar = 5 µm. (b) Shows chloroplasts (CL) in photosynthetic mesocarp cells followed by tube cells (TC), outer integument (OI), inner integument (II), and nucellar epidermis (NE). Thick deposits of cutin on the outer integument cell walls adjoining cross cells is with arrow heads. Bar = 10 µm. (c) Nucellar projection (NP) cells nearest the pigment strand (PS) have dense cytoplasm at 20 DPA. Bar = 5 µm. (d) Mitochondria is present in pigment strand cells whether dark deposits (DD) occur in those cells or not. Bar = 2 µm. (e) A section through adjacent inner integument cell and and a nucellar epidermis cell. Note the prominent plasmalemma between the cells. The cell wall of the nucellar epidermis cell has rich deposit of cellulose indicated with arrow heads. Bar = 2 µm. (f) At 30 DPA, the nucellar projection has dense cytoplasm and massive cell walls with frequent plasmodesmata connections (indicated with arrow heads). Bar = 5 µm.

### Pigment strand

The pigment strand is located inwards of the vascular bundle toward the endosperm. The pigment strand is remarkable for cells that accumulate dark deposits as the grain develops. At 5 DPA, the cells of the pigment strand can be distinguished from those of vascular parenchyma bundles by their elongate shape and generally denser cytoplasm. Dark deposits were detected on a few of the cells in the pigment strand at 5DPA. They appeared to first accumulate in small amount, in a subcellular organelle, possibly the vacuole. The organelle then enlarges, with the already accumulated deposit forming a continuous ring bordering the membrane of the organelle. The rest of the cytoplasm is pushed against the cell wall as the organelle expands. The initial clear space created within the organelle by its enlargement is eventually filled with the dark deposit as the grain matures (Fig. 2a-g; Fig. 3b-d). The number of pigment strand cells filled with dark deposit increased at subsequent sampling stages. Note that even at 30 DPA, there were still pigment strand cells that had not accumulated dark deposits. Abundant plasmodesmata connections were observed between adjacent pigment strand cells (whether filled with dark deposit or not), between pigment strand cells and vascular parenchyma cells, and, between pigment strand cells and nucellar projection cells (Fig. 3).

Four layers of different cell types flank the pigment strand on both sides and are continuous round the developing endosperm (Fig. 2a-b). They are tube cells (CC), outer integument (OI), inner integument (II) and nucellar epidermis (NE), in that order from the outer surface. For brevity we will refer to them as inner pericarp cells (IPC).

Tube cells are bordered on the outside by mesocarp cells and on the inside by the outer integument. They are easily distinguished from the mesocarp cells by their lack of chloroplasts, circular shape and sparse cytoplasm. Tube cells had thin cell walls in all the samples examined and enlarge as the grain develops till around 20 DPA. They subsequently appeared shrunk in size and separate from one another by 30 DPA.

The outer integument cells have sparse cytoplasm and enlarge as the grain develops till about 20 DPA. The cell walls of outer integument cells adjacent to tube cells are already cutinized at 5 DPA and progressively become heavily cutinized as the grain matures (Fig. 2b-g). The inner cell walls abutting the inner integument also become cutinized to a lesser degree as the grain develops. From 25 DPA, outer integument cells collapse and appear as one layer of cutin. The collapse of the outer integument cells starts from the dorsal grain surface and proceeds to the ventral surface.

Inner integument cells lie between the outer integument and nucellar epidermis. Inner integument cells are filled with dark deposits in the manner already described for pigment strand cells. The deposits have been identified in barley as tannins (Briggs, 1974; Felker et al., 1984).

Nucellar epidermis cells are the largest of the cell types in a developing Brachypodium grain at 5 DPA (Fig. 2a). The lateral nucellar epidermis cells are usually the largest (Opanowicz et al., 2011). Nucellar epidermis cells reach peak size at about 15 DPA. Subsequently, they gradually become smaller, possibly as a result of pressure from the expanding endosperm. Adjacent nucellar epidermis cells are separated by thin cell walls with prominent plasmodesmata.

However, nucellar epidermis cell walls in contact with endosperm cells and inner integument cells are greatly thickened (Fig. 3e). Gullion et al, (2012), detected (1-3) (1-4)-β-D glucan in the inner and outer cell walls of nucellar epidermis cells in 7 DPA developing Brachypodium grains. The cytoplasm of nucellar epidermis cells appeared vacuolated at early stages of development. At later stages of development the cells were found filled with electron lucent cellulosic material, previously identified as (1-3) (1-4)-b-D glucan (Guillon et al., 2012). A prominent middle lamella was observed between nucellar epidermis cells and inner integument cells (Fig. 3e).

There are plasmodesmata connections between adjacent pigment strand cells and cells of these four cell types on both flanks. Plasmodesmata connections were not observed between the different types of cells that make up the IPC.

### Nucellar projection

Nucellar projection cells adjoins the pigment strand inwards towards the endosperm. Brachypodium nucellar projection cells can be distinguished into two types; (a) smaller and circular type close to the pigment strand that have dense cytoplasm, (b) larger and elongate type close to the endosperm that have sparse cytoplasm (Fig. 2). The cell walls of nucellar projection cells greatly thicken as the grain matures, especially the elongate nucellar cells close to the endosperm (Fig. 3f). A progressive increase in the cell wall thickening of nucellar projection cells from the pigment strand to the endosperm was observed. Signs of lysing of nucellar projection cells close to the endosperm were observed on grain sections from 10 – 30 DPA.

Numerous plasmodesmata connections were observed between adjacent nucellar projection cells.

### Endosperm cavity

Endosperm cavity was not observed on Brachypodium grains sampled at 5 and 10 DPA. At 15 DPA (Fig. 2d), endosperm cavity was observed between the endosperms and nucellar projection. The cavity was still present at 20 DPA (Fig. 2e), though occupying a lesser area compared to 15 DPA. By 25 DPA, the cavity has been completely occluded by the expanding endosperm (Fig. 2f).

### Assimilate transfer pathway into developing Brachypodium grains

Delineation of assimilate transport pathway into developing Brachypodium grains was investigated using fluorescent dyes; symplastic tracer; 5-(6)-6-carboxyfluorescein diacetate (CFDA; Sigma) and apoplastic tracer; 1% Lucifer Yellow (LY; Sigma). The grains were incubated in the dye solutions for different lengths of time and the localization of the dyes were confirmed using fluorescence microscopy.

CFDA was readily observed on transverse free hand sections of the Brachypodium grains that had been incubated in the dye for 15min. CFDA fluorescence was distinctly observed on the vascular bundle, pigment strand cells, nucellar projection cell and round the grain in nucellar epidermis cells (Fig. 4). Similar, results were obtained on grains incubated in the dye for 30, 45 and 60min. As the period of incubation increased, dye fluorescence increased centripetally in the endosperm. This suggests that materials travel circumferentially around the grain and at some point cross into endosperm cells. So the longer the grains were incubated, the more CFDA is centripetally accumulated, evidenced by intense fluorescence in the endosperm of grains sectioned after 60min incubation in CFDA (data not shown).

**Fig. 4:**
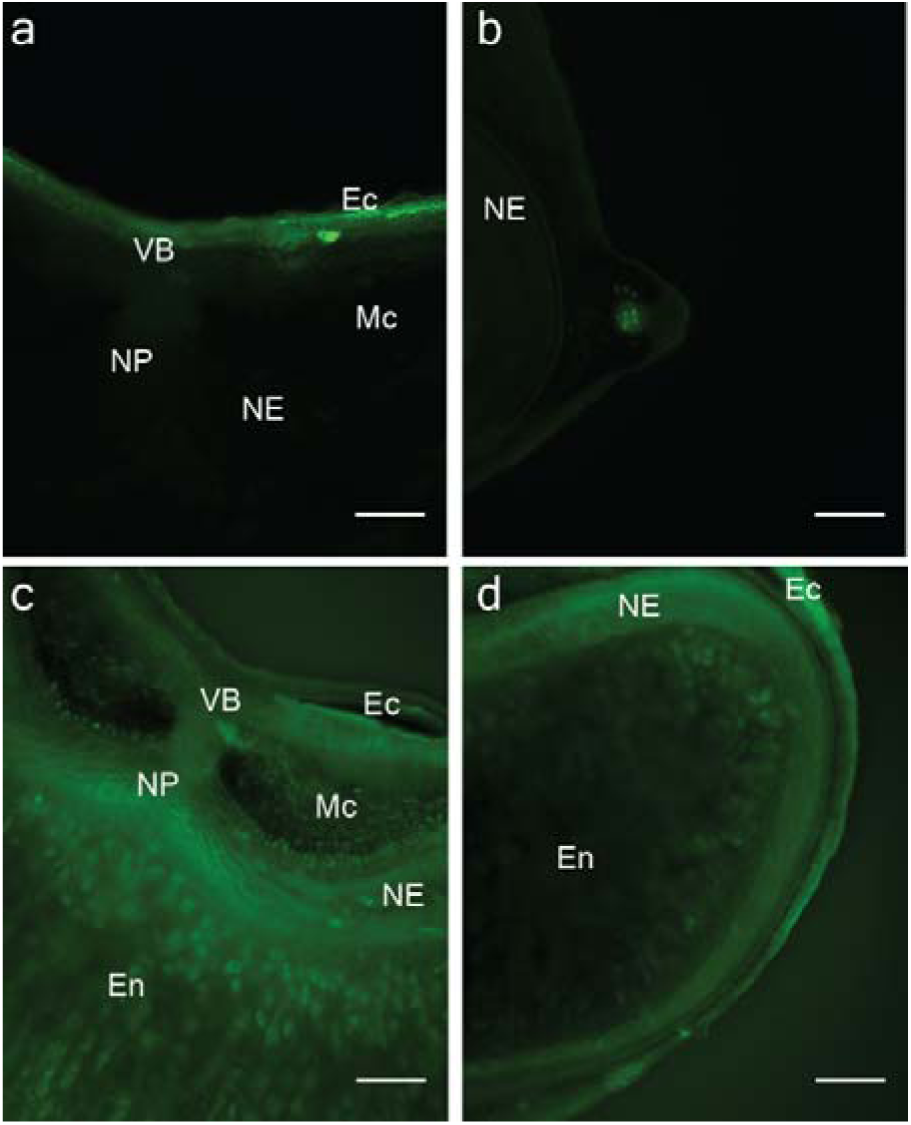
Delineation of assimilate transport route into Brachypodium endosperm using 5(6)-carboxyfluorescein diacetate (CFDA). (a), (b), (c), and (d) are free hand transverse section of 20 DPA Brachypodium grains. (a) and (b) shows ventral and lateral portions of sections that were incubated for 15 min in distilled water as control. There is auto fluorescence in the epicarp (Ec). (c) and (d) are similar portions of sections made from grains that were incubated in for 15 min in 0.01% CFDA. The dye signal is distinct in the vascular bundle (VB), nucellar projection (NP), nucellar epidermis (NE), and endosperm (En). Bar = 0.05 mm.

Fluorescent signals were not detected in sections of Brachypodium grains incubated in Lucifer Yellow for any of the period of time used in this experiment (data not shown). Since Lucifer yellow is an apoplastic dye, this suggests that assimilate movement into Brachypodium endosperm may be largely symplastic.

## Discussion

The succession of cell types in the post-phloem assimilate delivery pathway of Brachypodium represent a combination of features of such pathways reported in wheat, barley and rice. A ventral vascular bundle that spans the length of the caryopsis is a common feature in these three species (Thorne, 1985). The grain vascular bundle is connected to the rachilla and serves as route of water and assimilate transport from the rest of the plant into developing caryopsis (Frazier and Appalanaidu, 1965; Lingle and Chevalier, 1985). Our observation of the formation of semi-circle by the phloem of the vascular bundle of Brachypodium grains at 10 DPA (Fig. 2a, c) corresponds to observation of a phloem arc in rice caryopsis at 12 DPA (Oparka and Gates, 1981). The occurrence of immature sieve elements containing little cytoplasm alongside mature sieve elements with empty lumen has also been reported in rice (Oparka and Gates, 1981). Since the vascular bundle continue to differentiate as the grain develops, the immature sieve elements observed may be newly differentiated ones. Plasmodesmata connection among and between cells of the vascular bundle of Brachypodium grains suggest symplastic exchange of water and assimilates between these cells. This is similar to observations in rice and wheat (Oparka and Gates, 1981; H. L. Wang, C. E. Offler, et al., 1995). Also, Plasmodesmata connection between cells of the vascular bundle and surrounding parenchyma cells is consistent with observations in wheat (H. L. Wang, C. E. Offler, et al., 1995). This arrangement of cells suggest that assimilates move vertically up from the base of the grain through the vascular bundle and are discharged horizontally into the parenchyma cells along the way.

Sucrose is the main sugar transported in grasses (Thorne, 1985; van Bel and Hess, 2008). Sucrose is cleaved to hexoses on exit from the vascular bundle in maize and sorghum by vacuolar and cell wall invertases (Bihmidine et al., 2013; Felker and Shannon, 1980; Maness and McBee, 1986; Porter et al., 1985). In wheat and barley, whether or not sucrose is cleaved after it is unloaded from the vascular bundle remains a debate (Thorne, 1985). In any case, accumulation of glucose in wheat endosperm cavity sap coupled with 80% reduction of sucrose concentration compared to sieve tube sap strongly suggests that sucrose is cleaved in the post-phloem transfer cells of wheat (Fisher and Gifford, 1986). Whether or not sucrose is transformed in the post-phloem pathway of Brachypodium is an interesting research question. However, the dense cytoplasm of the vascular bundle and parenchyma cells suggest they are not just passive channels of for assimilate movement. The abundance of mitochondria in these cells suggest the occurrence of energy demanding processes. This sharply contrasts with surrounding mesocarp cells which gradually lose their cytoplasm as the grain develops, and but for chloroplasts, appear empty at 30 DPA.

The presence of pigment strand have been reported in barley rice and wheat (Felker et al., 1984; Oparka and Gates, 1982; H. L. Wang, C. E. Offler, et al., 1995). While the pigment strand cells of rice grain accumulate lipid deposits, those of wheat and barley accumulate phenolic deposits (Felker et al., 1984; Oparka and Gates, 1982). The pigment strand in Brachypodium is strikingly similar to barley in the extent it gets filled with dark deposits (Lingle and Chevalier, 1985). Dark deposits in Brachypodium pigment strand cells were visible at 5 DPA whereas they appears at about 12 DPA in barley (Felker et al., 1984; Lingle and Chevalier, 1985), suggesting early accumulation of dark deposits in Brachypodium. The accumulation of deposits in pigment strand cells was initially thought to limit assimilate transport towards the endosperm during wheat grain development (Zee and O’brien, 1970). Subsequent reports failed to confirm this view, but rather suggested that assimilate transport is restricted to the symplast at the pigment strand (Lingle and Chevalier, 1985; Oparka and Gates, 1982). On the basis of comparative anatomy, we speculate that assimilate movement into developing Brachypodium grains may not be limited by the accumulation of dark deposits in its pigment strand cells. Whether assimilate transport is restricted to the symplast at the pigment strand of Brachypodium cannot be firmly confirmed from our results. However, the high frequency of plasmodesmata connection between pigment strand cells and vascular parenchyma cells suggest an increased transportation capacity that may compensate for loss of apoplastic transport between the vascular parenchyma cells and the pigment strand cells.

Plasmodesmata connection between pigment strand cells and IPC, namely; tube cells, outer integument, inner integument and nucellar epidermis, suggest that assimilate supply to these cells may rely significantly on pigment strand cells. These cells are already well differentiated at 5 DPA. Our observations suggest little or no cell division in these four flanking cell types as the grain develops. They greatly enlarge and further differentiate by accumulation and synthesis of materials as during grain development. Since they lack chlorophyll, the materials they need are mostly likely supplied through the pigment strands. Thus, suggesting that Brachypodium pigment strand cells may serve as a hub for assimilate distribution to different types of cells. This therefore raises the question; to what extent does assimilate supply to other grain cell types deprive Brachypodium endosperm resources for starch synthesis? We plan to answer this question in a future experiment.

Based on cell morphology, we identified only two types of nucellar projection cells in Brachypodium, whereas three and four types of nucellar projection cells were identified in barley and wheat respectively (Thiel et al., 2008; Wang et al., 1994). However, as in barley and wheat, the nucellar projection cells closest to the pigment strand were the least differentiated and those closest to the endosperm had massive cell walls. Compared to wheat and barley nucellar projection cells do not undergo extensive autolysis as the grain develops in Brachypodium.

The limited lysing of nucellar projection cells (Fig. 2d-e) creates a transient cavity that is smaller but similar to the endosperm cavity of wheat and barley. The endosperm cavity serves as a temporary apoplastic sink for assimilates (H. L. Wang, C. E. Offler, et al., 1995; H. L. Wang, J. W. Patrick, et al., 1995). The presence of endosperm cavity contribute to grain shape in cereals (Hands et al., 2012). Thus, the transient endosperm cavity contribute to the narrow crease in a mature Brachypodium grains (Hands and Drea, 2012).

Whereas nucellar cells in barley, maize, rice, sorghum and wheat usually undergo programmed cell death (PCD) and lyse after fertilization (Radchuk and Borisjuk, 2014), Brachypodium nucellar cells are slow to die and the nucellar epidermis remain prominent and persistent throughout grain development (Solomon and Drea, unpubl. data). Remarkably, the cell walls of nucellar projection cells in developing Brachypodium grains become thick, but lack wall ingrowths as found in barley and wheat (Cochrane and Duffus, 1980; Wang et al., 1994). Wall ingrowths in nucellar projection cells and other types of transfer cells, potentially increase plasma membrane surface area for rapid assimilate transport (Offler et al., 2003; H. L. Wang, C. E. Offler, et al., 1995).

It is tempting to speculate that the lack of wall ingrowths in nucellar projection cells of Brachypodium grains may contribute to lower post-phloem assimilate transport capacity compared to wheat and barley. This speculation is plausible considering that assimilate delivery to developing barley and wheat endosperms relies only on nucellar projection cells for passage of assimilates from the pigment strand into the endosperm cavity. Hence the need for the increased surface area for rapid assimilate movement. Whereas in Brachypodium, assimilates are also transported through nucellar epidermis (Fig. 4), in addition to the nucellar projection. This additional channel for assimilate movement probably means that less assimilate pass through Brachypodium nucellar projection cells compared to wheat and barley. However, numerous plasmodesmata connection between cells of Brachypodium nucellar projection cells (despite thickened cell walls) suggest a capacity for rapid assimilate movement across these cells. The thick walls of Brachypodium nucellar projection cells may also provide apoplastic route of assimilate and water movement towards the endosperm as reported in wheat (H. L. Wang, C. E. Offler, et al., 1995).

Assimilate movement into Brachypodium endosperm is identical to rice where the main supply route is circumferentially via the nucellar epidermis (Fig. 1b). In rice, several sucrose transport genes have been shown to be expressed along the post-phloem assimilate delivery pathway to developing grains (Bai et al., 2016; Furbank et al., 2001; Ma et al., 2017). While some of the genes have active roles in sucrose export from the nucellar epidermis into the maternal/filial apoplast, others have been linked with assimilate import into aleurone cells (Ma et al., 2017). Mutation of sucrose transport genes and/or their regulators negatively affects grain filling and lead to abnormally shaped rice grains (Bai et al., 2016; Ma et al., 2017; Wu et al., 2018). Though sucrose transport genes are conserved in Brachypodium (Braun and Slewinski, 2009), their role in assimilate transport and grain filling is yet to be reported. Nevertheless, a major structural difference is that in rice, nucellar epidermis cells collapse during grain development while in Brachypodium they persist till grain maturity. Based on earlier histological studies, it was suggested that mechanical compression of rice nucellar epidermis by expanding endosperm prevents further flow of assimilates and terminate grain filling (Ellis and Chaffey, 1987; Ellis et al., 1987). However, recent findings have demonstrated programmed cell death (PCD) in nucellar tissues of developing rice grains (Yang et al., 2012; Yin and Xue, 2012). It has also been shown that perturbation of genes involved in PCD of nucellar tissues adversely affects grain filling and lead to abnormally shaped grains (Nayar et al., 2013; Yang et al., 2012; Yin and Xue, 2012). Furthermore, PCD of nucellar tissues is known to be generally necessary for normal grain development in cereals (Domínguez and Cejudo, 2014; Lu and Magnani 2018). It is therefore remarkable that normal grain development and filling in Brachypodium does not require disintegration of nucellar epidermis cells. We previously showed that nucellar epidermis cells are dead in mature Brachypodium grain (Hands et al., 2012). However, there is no data yet on when and how its death affects grain development and filling. An additional feature of Brachypodium nucellar epidermis cells is accumulation (1-3) (1-4)-β-D glucan as the grain develops (Guillon et al., 2012). It is not clear if and how accumulation of (1-3) (1-4)-β- D glucan in nucellar epidermis cells affect assimilate transport through these cells. It is possible that (1-3) (1-4)-β-D glucan accumulation in the nucellar epidermis cells deprive the endosperm of assimilates and decrease the rate of assimilate transport.

## Conclusion

The fine structure and post-phloem assimilate transport into developing grains of Brachypodium reflects the species phylogenetic position between Ehrhartoideae and Pooideae. Brachypodium post-phloem assimilate delivery pathway is structurally similar to wheat and barley save for the absence of modified aleurone. Its assimilate delivery strategy on the other hand is identical rice. These combination of temperate anatomy with tropical physiology marks Brachypodium as an excellent model to understand the evolution of specialised assimilate transfer cells (modified aleurone) for efficient assimilate acquisition in temperate cereal grains. Such knowledge can be exploited in domestication of new crop species.

## Abbreviations

bd: basal direct
CEAAR: Caryopsis Endosperm Assimilate Acquisition Route
DPA: Days Post Anthesis
IPC: Inner Pericarp Cells
vc: ventral circuituous
vd: ventral direct

## Acknowledgements

We are grateful to Natalie Allcock and Anna Straatman-Iwanowska for assistance with grain sectioning and electron microscopy. CUS is a PhD student funded by Tertiary Education Trust Fund, Nigeria.

